# The Burden Borne by Protein Methyltransferases: Rates and Equilibria of Nonenzymatic Methylation of Amino Acid Side-Chains by SAM in Water

**DOI:** 10.1101/2021.03.17.435195

**Authors:** Charles A. Lewis, Richard Wolfenden

## Abstract

SAM is a powerful methylating agent, with a methyl group transfer potential matching the phosphoryl group transfer potential of ATP. SAM-dependent N-methyltransferases have evolved to catalyze the modification of specific lysine residues in histones and transcription factors, in addition to generating epinephrine, N-methylnicotinamide, and a quaternary amine (betaine) that is used to maintain osmotic pressure in plants and halophilic bacteria. To assess the catalytic power of these enzymes and their potential susceptibility to transition state and multisubstrate analogue inhibitors, we determined the rates and positions of equilibrium of methyl transfer from the trimethylsulfonium ion to model amines in the absence of a catalyst. Unlike the methyl group transfer potential of SAM, which becomes more negative with increasing pH throughout the normal pH range, equilibrium constants for the hydrolytic demethylation of secondary, tertiary and quaternary amines are found to be insensitive to changing pH and resemble to each other in magnitude, with an average ΔG value of ∼ -0.7 kcal/mol at pH 7. Thus, each of the three steps in the mono- di- and trimethylation of lysine by SAM is accompanied by a free energy change of -7.5 kcal/mol in neutral solution. Arrhenius analysis of the uncatalyzed reactions shows that the unprotonated form of glycine attacks the trimethylsulfonium ion (TMS+^+^) with a second order rates constant of 1.8 × 10^−7^ M^-1^ s^-1^ at 25 °C (ΔH^‡^ = 22 kcal/mol and TΔS^‡^ = -6 kcal/mol). Comparable values are observed for the methylation of secondary and tertiary amines, with k_25_ = 1.1 × 10^−7^ M^-1^ s^-1^ for sarcosine and 4.3 × 10^−8^ M^-1^ s^-1^ for dimethylglycine. The nonenzymatic methylation of imidazole and methionine by TMS+^+^, benchmarks for the methylation of histidine and methionine residues by SETD3, exhibit k_25_ values of 3.3 × 10^−9^ and 1.2 × 10^−9^ M^-1^ s^-1^ respectively. Lysine methylation by SAM, although slow under physiological conditions (t_1/2_ 7 weeks at 25 °C), is accelerated 1.1 × 10^12^ -fold at the active site of a SET domain methyltransferase.

---

Histones and transcription factors are equipped with lysine residues whose ε-amino groups are subject to enzymatic mono-, di- or trimethylation in the presence of highly specific SAM^+^-dependent N-methyltransferases (KMTs), a central event in the regulation of gene transcription in eukaryotic organisms. The crystal structures of KMTs with analogous “SET” domains (the acronym denotes “Su(var), Enhancer of zeste, Trithorax”) reveal the presence of individual substrate binding sites for SAM^+^ and the methyl-accepting nitrogen atom of the acceptor. These sites are separated by a pore in the enzyme structure through which a methyl group is transferred from SAM at each stage in the mono-, di- and trimethylation of the ε-amino group of lysine^1-3^ Other SAM-dependent N-methyltransferases are responsible for generating quaternized amines such as N,N,N-trimethylglycine (betaine) that maintain osmotic pressure in halophilic bacteria and plant cells,^4^ and for the production of epinephrine and N methylnicotinamide, and powerful inhibitors of these enzymes have been obtained by combining the binding determinants of SAM and the methylation target within a single stable molecule.^*5-7*^

To evaluate the rate enhancements that these enzymes produce—and their potential susceptibility to inhibition by transition state or multisubstrate analogues—it would be useful to have information about the corresponding reactions in the absence of enzymes. Uncatalyzed reaction rates have been reported for other SAM-dependent methyl transfer reactions, including those catalyzed by the “halogenases” that employ halide ions to generate atmospheric chloromethane and fluoromethane from seawater;^8^ and for “etherases” that are responsible for recycling much of the carbon in the biosphere.^9^ But apart from a single data point in the literature,^10^ methyl transfer to amines appears to have attracted little attention. For comparison with SET domain methyltransferases, it would be useful to know how much primary, secondary and tertiary amines—which vary over 15 orders of magnitude in their intrinsic basicities in the vapor phase^11^—vary in their susceptibility to methylation by sulfonium ions in water. SETD3, which is similar in structure to SET domain KMTs and is weakly active on lysine, also catalyzes the methylation of histidine and methionine side-chains in actin.^12^

To obtain benchmarks for these activities, we conducted experiments at elevated temperatures to determine the rates and equilibria of nonenzymatic methylation of aliphatic primary, secondary and tertiary amines, imidazole and methionine.

## RESULTS AND DISCUSSION

### Equilibria of methyl transfer between amines, sulfonium ions and water

In preliminary experiments, we sought to determine the equilibrium constant for hydrolytic cleavage of methanol (MeOH) from each of the methylamines indirectly, by measuring the equilibrium constant for methyl transfer to each amine from TMS^+^ whose equilibrium constant for hydrolysis is known.^13^ At 200 °C, the tetrafluoroborate salt of the trimethylsulfonium ion (TMS^+^) reacted with methylamine (MeNH_2,_ half-titrated with HCl) to yield reaction mixtures containing di-, tri- and tetramethylamine. However, the progressive hydrolysis of the sulfonium ion—occurring in competition with the methylation of MeNH_2_—generated sufficient acid to reduce the pH continuously, interfering with accurate determination of the position of the equilibrium of the reaction, which releases a proton (equation 1).

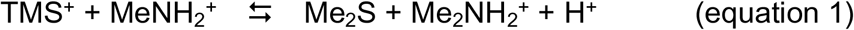

To circumvent that difficulty, we adopted a more direct approach, heating MeNH_2_ (0.55 M) with MeOH (1.1 M) in 1 M HCl, in quartz tubes sealed under vacuum. After various time intervals, reactions were stopped with ice and the product mixtures were analyzed by proton NMR. After 120 h at 250 °C, we observed the appearance of di-, tri- and tetramethylammonium ions at concentrations that arrived at a constant ratio: [MeOH] = 0.096 M; [MeNH_3_ ^+^] = 0.296 M; [Me_2_ NH_2_ ^+^] = 0.112 M; [Me_3_ NH^+^] = 0.027 M; [Me_4_N^+^] 0.0049 M. Under these conditions, the estimated half-life for approach to equilibrium was ∼20 h. The relative concentrations of these species showed little variation when the same experiment was conducted over the temperature range from 240 to 290 °C (Figure 1), and yielded the following equilibrium constants for each of the three steps in the hydrolytic demethylation of the tetramethylammonium ion, taking the activity of water as unity by convention (equations 2-4):

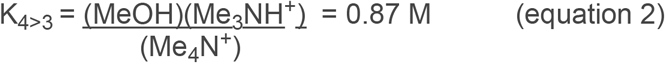

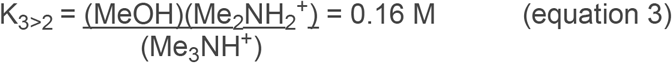

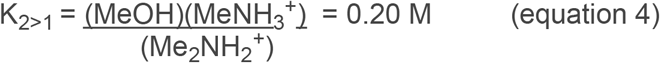

**Figure 1:**
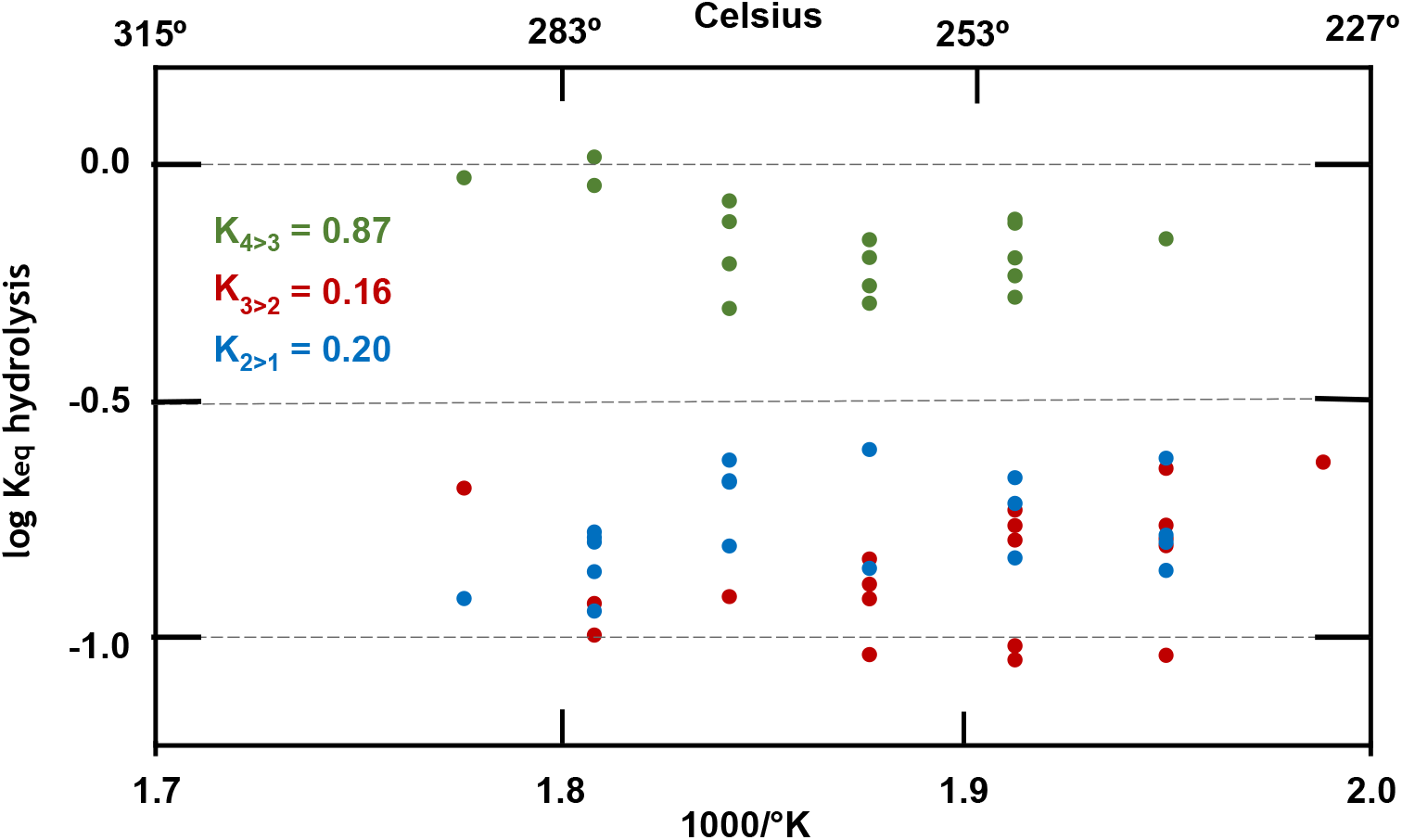
van’t Hoff plot of equilibrium constants for the hydrolytic demethylation of mono-, di- and trimethylamine (equations 2-4), plotted as a logarithmic function of the reciprocal of absolute temperature.

Separate experiments, performed with NH_4_Cl (1 M) and MeOH (1 M) in 1 M HCl, yielded a slightly smaller equilibrium constant for the hydrolytic deamination of MeNH_3_^+^ (equation 5).

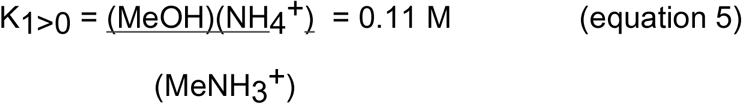

Equation 4, with an average value (K_2>1_) of 0.20 M at 250 °C, is analogous to the hydrolytic cleavage of a methyl group from the monomethylated ε-amino group of a lysine side-chain. To provide a basis for extrapolation to 25 °C, we used standard bond energies (or enthalpies) ^14^to estimate the value of ΔH for dimethylamine hydrolysis in the vapor phase, obtaining a value of -5.1 kcal/mol. Adding a small correction for the change in enthalpy of solvation that accompanies hydrolysis (+0.7 kcal/mol), based on Cabani’s tables,^15^ resulted in a total enthalpy change of -4.4 kcal/mol for hydrolysis of Me_2_NH_2_ ^+^. That value corresponds to a 24.7-fold increase in K_2>1_, from a value of 0.20 M measured at 250 °C to an estimated value of 4.9 M at 25 °C. Accordingly, our best estimate of the methyl group transfer potential (ΔG^0’^_pH 7_) of Me_2_NH_2_ ^+^ (or, by extension, the side-chain of an ε-monomethylated lysine residue in a solvent-exposed position within a protein) at 25 °C is -0.7 kcal/mol.

The pK_a_ values of methylamine and dimethylamine are virtually identical (10.64 and 10.67, respectively).^16^ The near-equivalence of these values (K_a(1)_ and K_a(2)_ in Scheme 1) implies that the affinities of amines and ammonium ions for methyl groups, as measured by the hydrolytic equilibrium constants K_1_ and K_2_ in Scheme 1, must also be virtually identical. For that reason, the free energy of hydrolysis of the N-CH_3_ bond of an ε-N-methylated lysine residue is not expected to vary significantly over the pH range from 0 to 14 (Scheme 2, blue line) and the values of K_2>1_ and K_3>2_—evaluated for each of the methylammonium ions in the present experiments—should apply to the uncharged amines as well. In contrast, ΔG^0’^ for hydrolytic demethylation of SAM^+^ becomes increasingly negative throughout the pH range from -5 to +14, reaching a value of -8.2 kcal/mol in neutral solution.^13^ Accordingly, the driving force behind the initial methylation of lysine by SAM^+^ is -7.5 kcal/mol in neutral solution (Scheme 2). Moreover, the similarity of the individual equilibrium constants associated with equations 2-4 (Figure 1) implies that the equilibrium constants for di- and trimethylation of lysine residues by SAM^+^ must also be similar.

**Scheme 1.**
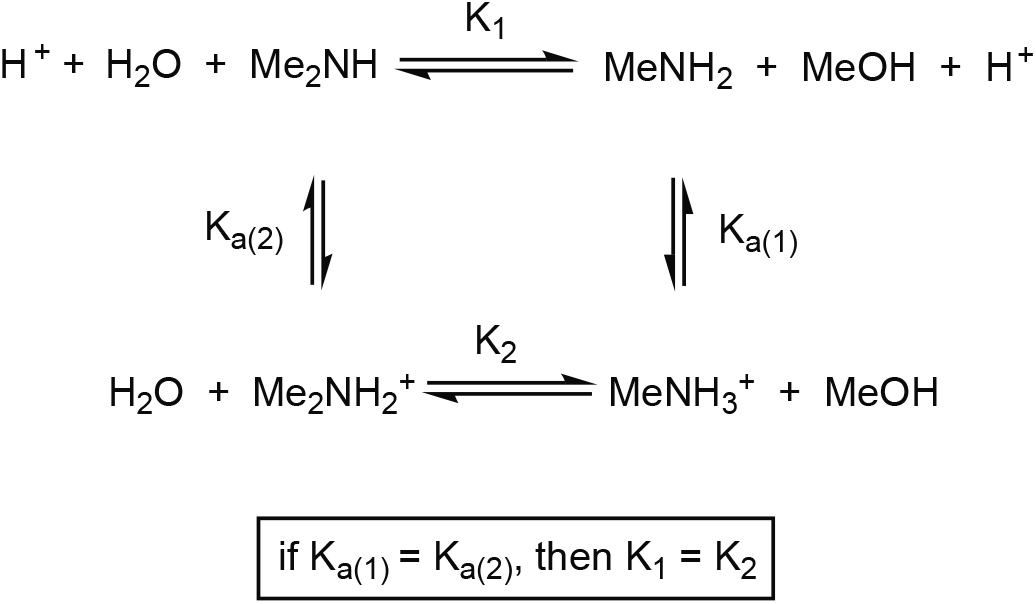
Relationship between the K_a_ values of Me_2_NH_2_^+^ (K_a(2)_) and MeNH_3_^+^ (K_a(1)_) and the equilibria of hydrolytic demethylation of Me_2_NH_2_^+^ (K_2_) and Me_2_NH (K_1_).

**Scheme 2.**
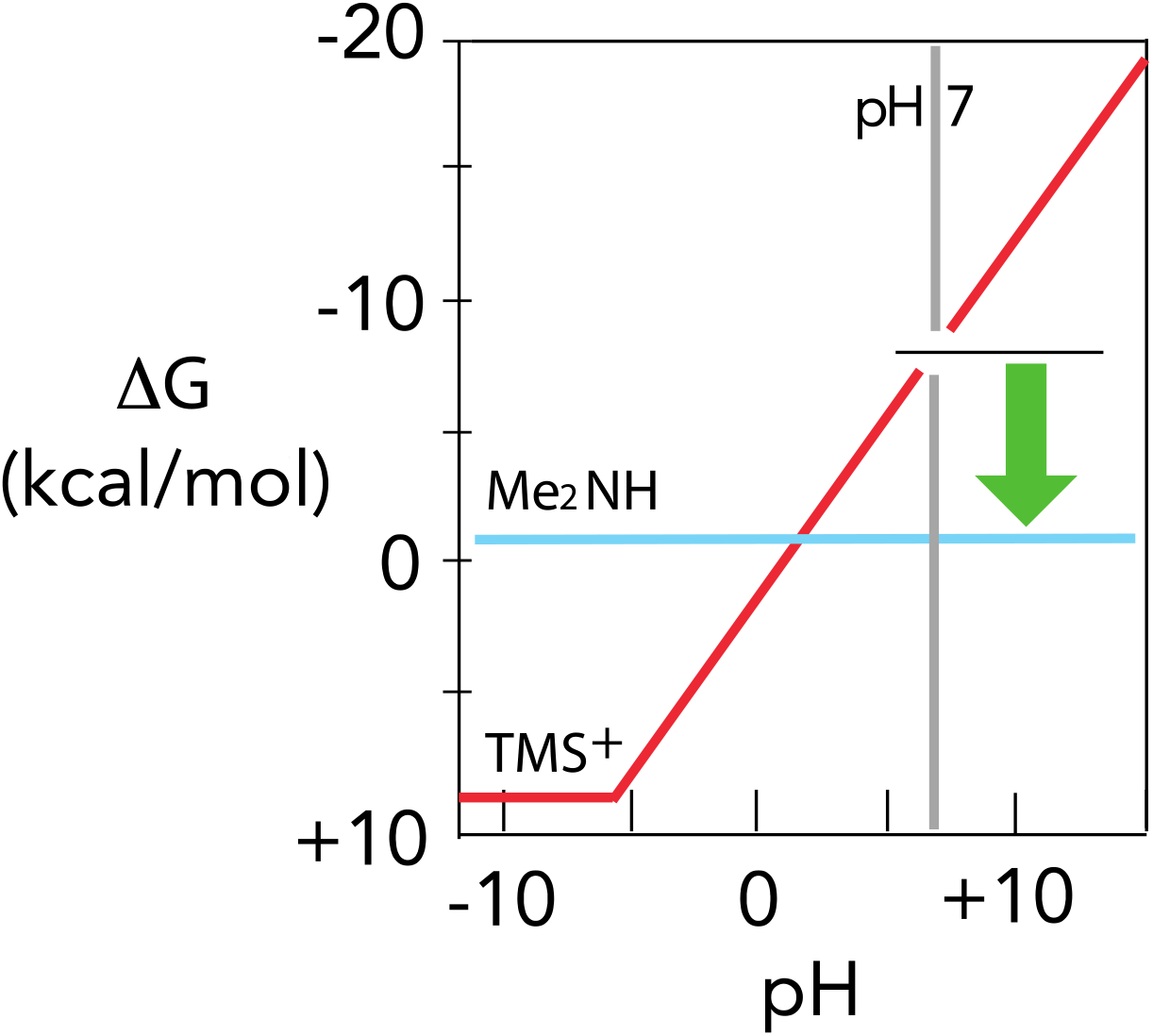
Influence of pH on the free energies of hydrolysis of SAM (as approximated by TMS^+^) (red line) and of Δ-N-methyllysine (as approximated by the hydrolytic demethylation of Me_2_NH) (blue line).^13^ Since the pK_a_ values of MeNH_3_^+^ and Me_2_NH_2_^+^ are similar in water, the position of equilibrium for the hydrolytic demethylation of Me_2_NH_2_^+^ is expected to be insensitive to changing pH (Scheme 1). In contrast, the free energy of methyl transfer from TMS^+^ to MeNH_2_ is strictly dependent on pH, arriving at an estimated value of -7.5 kcal/mol at pH 7 (green arrow).

### Rates of uncatalyzed methylation of the side-chains of lysine, histidine and methionine by sulfonium ions

In kinetic experiments to model the enzyme-catalyzed trimethylation of an exposed histone lysine side-chain by SAM—the process catalyzed by KMTs—reaction mixtures typically contained TMS^+^:BF^4-^(0.1 M), the amine hydrochloride (0.5 M) and KOH (0.25). Sealed in quartz tubes under vacuum, samples were incubated at temperatures ranging from 40 to 120 °C in ovens equipped with ASTM thermometers. At intervals, samples were removed and analyzed by ^1^H-NMR after exact dilution with D_2_O to which pyrazine had been added as an integration standard. In each case, the decomposition of TMS^+^ followed simple first-order kinetics for at least 3 half-lives. In the case of the primary amines methylamine and glycine, the chief product (in addition to dimethylsulfide which is volatile at ordinary temperatures) was the singly methylated starting material, along with small concentrations (<4% in total) of doubly and triply methylated amine, and methanol. In these experiments, the extent of hydrolysis of TMS^+^—as indicated by the appearance of methanol—was never greater than 1%. To obtain the second order rate constant (k_2_) for each methylation reaction, the first order rate constant for disappearance of TMS^+^ (k_1_) was divided by the starting concentration of unprotonated primary, secondary or tertiary amine and multiplied by the fraction of each of those nucleophiles appearing as its singly methylated product. To minimize the effects of acid generated during these reactions (equation 1), in which each reactant amine served as its own buffer, calculated rate constants were based on observations during the first 5% of reaction. The sensitivity and reproducibility of the integrated intensities of the reactant and product peaks allowed rate constants to be determined with an estimated error of ±5% in all cases. The results of these experiments yielded linear Arrhenius plots of log_10_ k_2_ versus 1000/T that were used to estimate the rate constant for amine attack on TMS^+^, extrapolated to 25 °C. Values of ΔH^‡^ were obtained from the slopes of these Arrhenius plots, and values of TΔS^‡^ were obtained by subtracting ΔG^‡^ from ΔH^‡^. Estimated errors are: ΔG^‡^ = ± 0.4 kcal/mol, ΔH^‡^ = ± 0.7 kcal/mol, TΔS^‡^ = ± 1.0 kcal/mol. Similar procedures were used to examine the reactions of TMS^+^ with imidazole and methionine, chosen to represent the side-chains of His and Met residues in actin, which have recently been shown to undergo methylation in the presence of SETD3.^12^ We also examined the reaction of pyridine with TMS^+^ as a model for the enzymatic methylation of nicotinamide.^.5,6^ The results are summarized in Table 1 and Scheme 3.

**Table 1:**
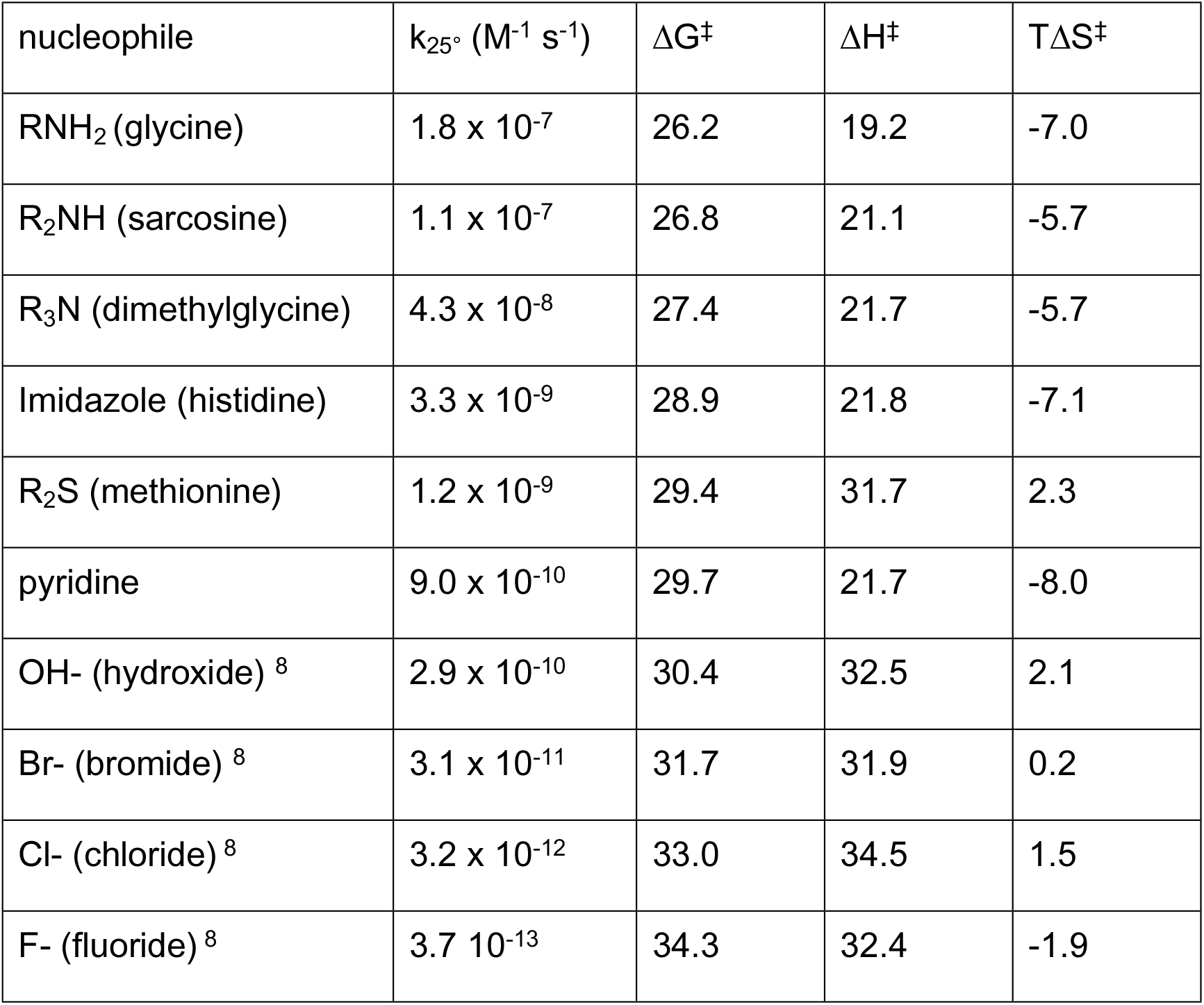
Rate constants and thermodynamics of activation (kcal/mol) for the nonenzymatic reactions of TMS^+^ with glycine, sarcosine, dimethylgycine, imidazole, pyridine and the sulfur atom of methionine (present work) compared with the reactions of TMS^+^ with hydroxide and halide ions (ref. 8). Estimated errors are: ΔG^‡^ = ± 0.4 kcal/mol, ΔH^‡^ = ± 0.7 kcal/mol, TΔS^‡^ = ± 1.0 kcal/mol.

**Scheme 3.**
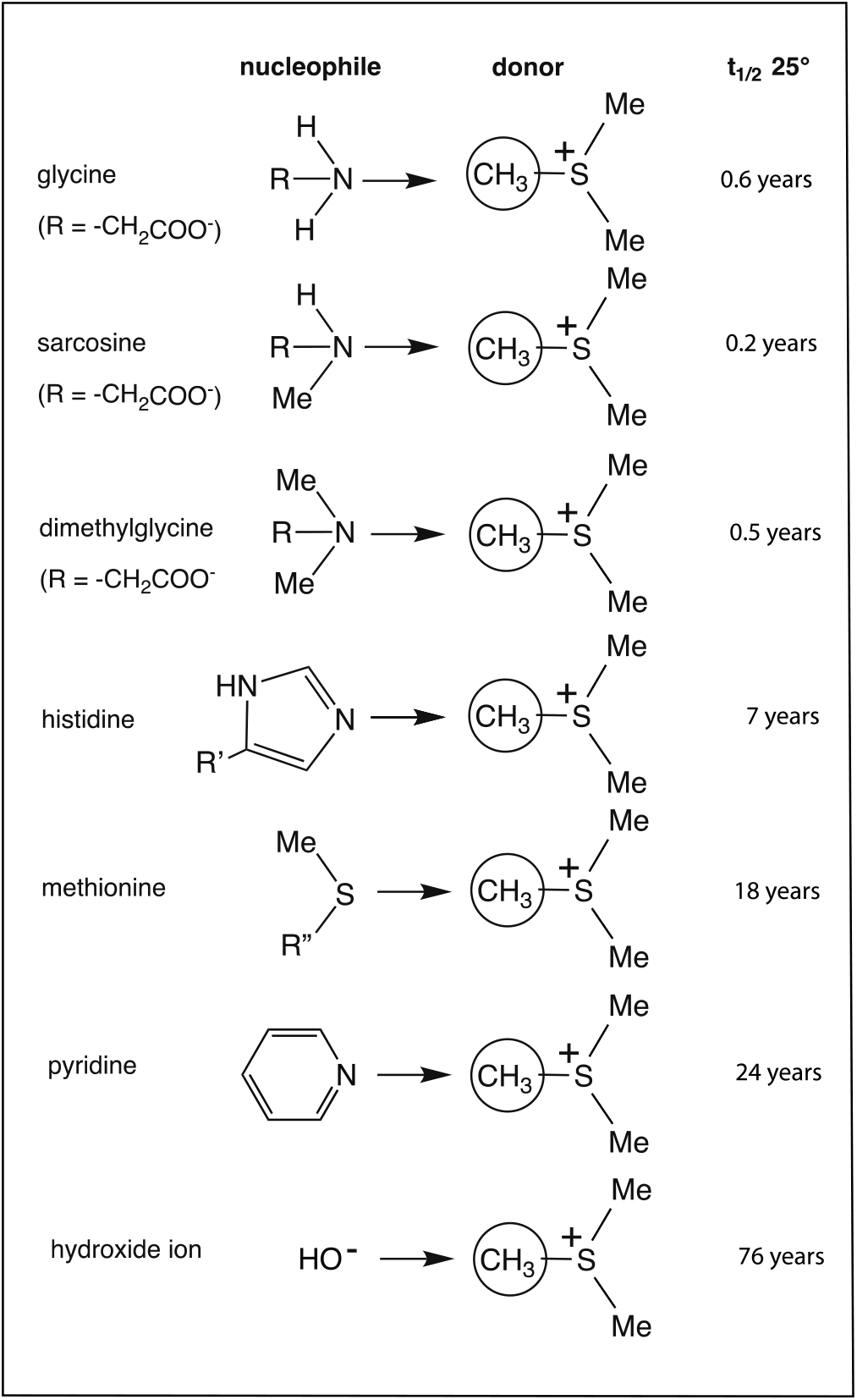
Half-lives for attack on TMS^+^ by model nucleophiles representing amino acid side-chains and pyridine. At the pK_a_ values of their conjugate acids, the rate of TMS^+^ hydrolysis is negligible, as indicated by the rate of attack by the hydroxide ion (ref. 8) which has been included for comparative purposes.

From a kinetic standpoint, the nonenzymatic methylation of amines proceeds much more rapidly than the methylation of halide ions or the S-methylation of methionine; and Mihel et al. have estimated an even lower rate constant (1.7 × 10^−14^ M^-1^ s^-1^ at 37 °C) for the uncatalyzed methylation of catechol by SAM^+^.^17^ In agreement with their observations, we were unable to detect significant methylation of phenolate or catecholate ions within the time required for complete hydrolysis of TMS^+^ at elevated temperatures.

### Comparison of the catalytic power and transition state affinity of lysine N-methyltransferases with those of other N-methyltransferases and two-substrate enzymes

The rate enhancement produced by the Rubisco large subunit KMT can be estimated by dividing the second order rate constant (k_cat_/K_m_) for this two-substrate enzyme reaction—in the presence of saturating SAM—by the present second order rate constant for reaction of a primary amine with TMS^+^. In attempting that comparison, it is worth noting that TMS^+^ is an imperfect model for SAM^+^, which can decompose by several routes^18^ and whose reactivity as a methylating agent may be depressed or enhanced by the presence of its homoseryl and adenosyl substituents. The three indistinguishable methyl groups of TMS^+^ would be expected to render it 3-fold more reactive than SAM^+^ as a methylating agent, and that statistical effect should be taken into consideration in using it as a model for SAM^+^.^8^ Trievel et al. report a K_m_ value of 1.4 × 10^−6^ M and a k_cat_ value of 0.047 s^-1^ for the Rubisco large subunit N-methyltransferase from peas.^1^ Comparison of the resulting k_cat_/K_m_ value (3.4 × 10^4^ M^-1^ s^-1^) with the present second order rate constant for the nonenzymatic reaction of MeNH_2_ with TMS^+^ at 25 °C (1.1 × 10^−7^ s^-1^ M^-1^), after a 3-fold reduction of the latter value to take account of the statistical effect mentioned above, indicates that the Rubisco KMT produces a rate enhancement of 1.1 × 10^12^ -fold. A similar k_cat_/K_m_ value (1.5 × 10^4^ s^-1^ M^-1^) has been reported for human phenethanolamine N-methyltransferase acting on trifluoroethanolamine,^19^ implying a similar second order rate enhancement of 4.1 × 10^11^- fold. Combined with the K_m_ value (4.7 × 10^−6^ M)^1^ of Rubisco KMT for the second substrate, SAM,^1^ these rate enhancements indicate that the nominal affinities of Rubisco KMT for a multisubstrate analogue inhibitor incorporating the binding affinities of both substrates in the transition state for direct methyl transfer would correspond to a K_d_ value of 5.0 × 10^−18^ M.^20^ Because binding by KMT would seem to require threading a “dumbbell-shaped” inhibitor through the narrow pore between the methyl donor and acceptor binding sites of Rubisco KMT, affinities of that magnitude may not be easy to realize in an actual inhibitor.

It is also of interest to compare the rate enhancement produced by the Rubisco large subunit N-methyltransferase with those produced by other two-substrate enzymes involved in group transfer (**Table 2**). With a second order rate constant of 1.1 × 10^−7^ s^-1^ M^-1^ at 25 °C, the rate of uncatalyzed methyl transfer from 1 M TMS^+^ to primary amines corresponds to a half-life of 2 months and, as noted above, an N-methyltransferase rate enhancement of 1.1 × 10^12^-fold. With a second order rate constant of 3.9 × 10^−9^ s^-1^ M^-1^, the rate of uncatalyzed phosphoryl transfer from 1 M ATP to glucose corresponds to a half-life of 6 years and a hexokinase rate enhancement of 5 × 10^14^-fold.^21^ With a second order rate constant of 6 × 10^−11^ s^-1^ M^-1^, the rate of the uncatalyzed reaction of 1 M glutathione with lignin corresponds to a half-life of 370 years and an etherase rate enhancement of 2 × 10^15^-fold.^9^ With an estimated second order rate constant of 5 × 10^−12^ s^-1^ M^-1^, the rate of the uncatalyzed reaction of 1 M lactate with NAD corresponds to a half-life of 4400 years and a lactate dehydrogenase rate enhancement of 2 × 10^15^-fold.^22^ With a second order rate constant of 3.7 × 10^−13^ s^-1^ M^-1^, the rate of uncatalyzed methyl transfer from 1 M SAM to the chloride ion corresponds to a half-life of 60,000 years and a chlorinase rate enhancement of 1.2 × 10^17^-fold.^8^ Of the two-substrate enzymes that have been characterized in this way, only the peptidyl-transferase center of the ribosome is known to produce a smaller rate enhancement. With a second order rate constant of 3 × 10^−5^ s^-1^ M^-1^, the rate of uncatalyzed transfer of fluorophenylalanine from its trifluoroethyl ester to 1 M glycinamide corresponds to a half-life of 6.4 hours and a rate enhancement 2.5 × 10^7^-fold.^23^

**Table 2:**
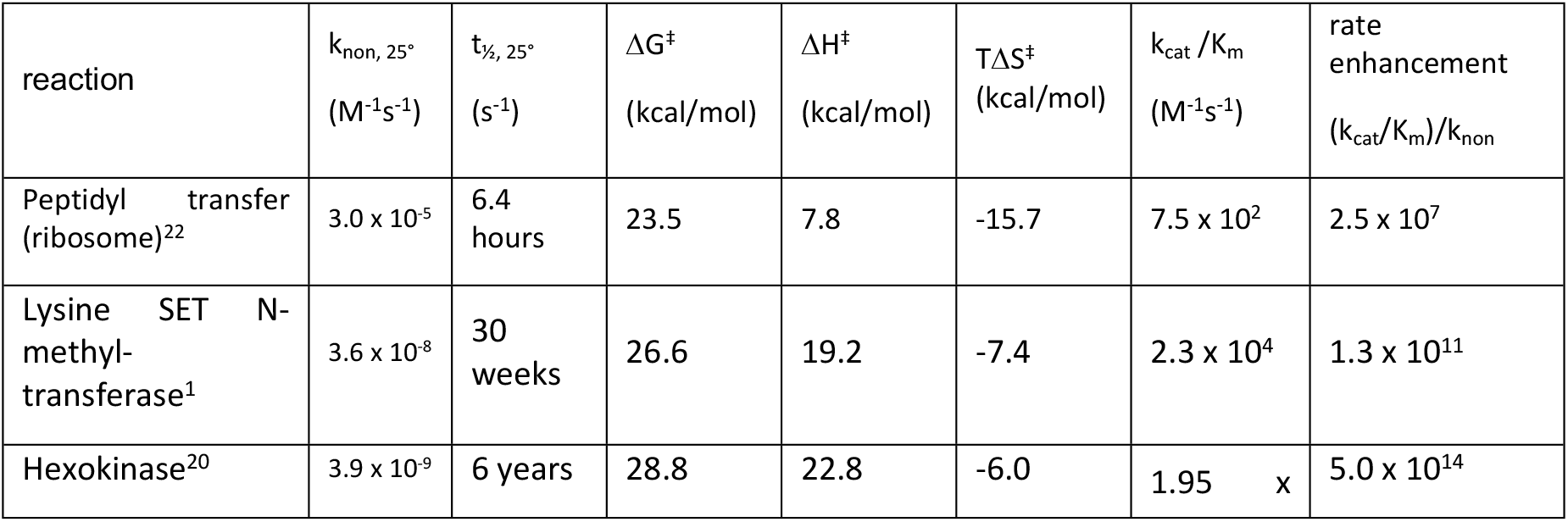
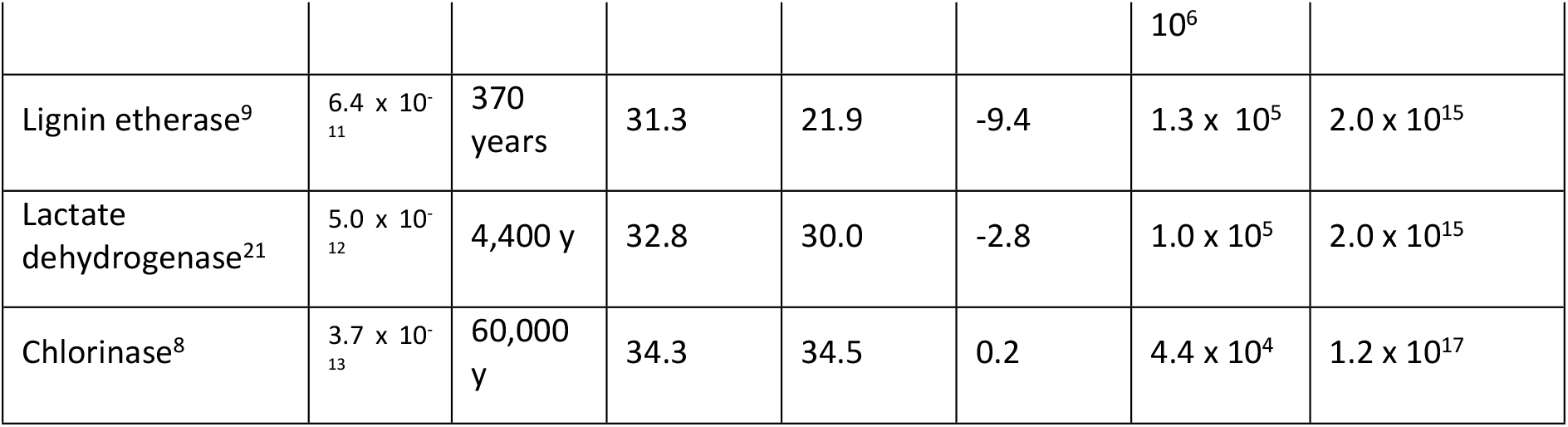
Second order rate constants and thermodynamics of activation for 2-substrate reactions in the absence of a catalyst, compared with k_cat_/K_m_ for the corresponding enzymes.

As is the case for many single-substrate and hydrolytic enzymes,^24,25^ variations in the magnitudes of these two-substrate rate enhancements can be seen in **Table 2** to arise largely from variations in the rate of the uncatalyzed reaction (k_non_) which, in the case of amine methylation, proceeds with a half-life (t_1/2_) of ∼ 7 weeks in the presence of 1 M SAM at 25 °C. At those rates, and with SAM present at much lower concentrations, the present nonenzymatic reactions seem unlikely to pose a threat to cell survival and reproduction under ordinary conditions. Moreover, Paik et al. have shown that several proteins undergo slow methylation of Asp and Glu residues by SAM in neutral solution the absence of enzymes, but detected no significant incorporation of alkali-stable methyl groups whose presence might have signaled N-methylation of Lys or Arg.^26^

In neutral solution—in the absence of a catalyst—the rates of the present reactions (Table 1), like their equilibrium constants (equations 2-5), are similar at each stage of methylation. The existing crystal structures of these N-methyltransferases^1-3^ appear to be consistent with a simple in-line displacement reaction. Moreover, the detailed structure of the lysine-binding cleft has also been shown to account for differences in specificity, between a SET domain KMT which catalyzes the mono-, di- and trimethylation of lysine,^1,2^ and the human KMT SET 7/9 which catalyzes only monomethylation and cannot accommodate methyllysine in a position appropriate for a second methyl transfer.^2^ Participation of a conserved tyrosine residue, acting as a general base, seems to have been ruled out by observations on a transition state – like complex,^3^ and would be unlikely to be helpful at the high pH values where these enzymes are active.^3^ Hughes and Ingold showed that nucleophilic displacement reactions—in which electrostatic charge becomes delocalized in the transition state— can be greatly accelerated when the substrates are removed from water.^27^ Binding determinants at the active sites of N-methylases, like those of etherases and halogenases, probably achieve much of their catalytic effect by relieving the substrates from constraints on their reactivity that were imposed by solvating water molecules before binding occurred.

## Footnotes

1 Wolfenden, R. (1972) Analog Approaches to the Structure of the Transition State in Enzyme Reactions, *Accts. Chem. Res. 5*, 10-18.

## REFERENCES

(1) Trievel, R. C., Beach, B. M., Dirk, L. M., Houtz, R. L., Hurley, J. H. (2002) Structure and Catalytic Mechanism of a SET Domain Methyltransferase. Cell 111, 91–103.

(2) Xiao, B. Jing, C., Wilson, J. R., Walker, P. A., Vasisht, N. Kelly, G., Howell, S., Taylor, I. A., Blackburn, G. M., Gamblin. S.J. (2003) Structure and Catalytic Mechanism of the Human Histone Methyltransferase SET 7/9, Nature 421, 652–656.

(3) Trievel, R.C., Flynn, E. M, Houtz, R. L., Hurley, J. H. (2003) Mechanism of Multiple Lysine Methylation by the SET Domain Enzyme Rubisco LSMT Nature Structural Biology 10, 545–552.

(4) Chen, S-Y., Lai, M-C., Lai, S-J. (2009) Characterization of Osmolyte Betaine Synthesizing Sarcosine Dimethylglycine N-Methyltransferase from Methanohalophilus portucalensis. Arch. Microbiol. 191, 735–743.

(5) Policarpo, R. L., Decultot, L., May, E., Kuzmic, P., Carlson, S., Huang, D., Chu, V., Wright, B. A., Dhakshinamoorthy, S., Kannt, A., Rani, S., Dittakavi, S., Panarese, J. D., Gaudet, R., Shair, M. D. (2019) High-Affinity Alkynyl Bisubstrate Inhibitors of Nicotinamide N-Methyltransferase (NNMT). J. Med. Chem. 62, 9837−9873.

(6) Chen, D., Li, L., Diaz, K., Iyamu, I. D., Yadav, R., Noinaj, N., Huang, R. (2019) Novel Propargyl-Linked Bisubstrate Analogues as Tight-Binding Inhibitors for Nicotinamide N-Methyltransferase. J. Med. Chem. 62, 10783−10797.

(7) Mahmoodi, N., Harijan, R. K., Schramm, V. L. (2020) Transition-State Analogues of Phenylethanolamine N-Methyltransferase. J. Am. Chem. Soc. 142, 14222−14233,

(8) Lohman, D. C., Edwards, D. R., Wolfenden, R. (2013) Catalysis by Desolvation: The Catalytic Prowess of SAM-dependent Halide-alkylating Enzymes. J. Am Chem. Soc. 135, 14473–14475.

(9) Lewis, C. A. Jr.,, Wolfenden, R. (2019) Ether Hydrolysis, Ether Thiolysis and the Catalytic Power of Etherases in the Disassembly of Lignin. Biochemistry 53, 5381–5385.

(10) Callahan, B. P., Wolfenden, R. (2003) Migration of Methyl Groups Between Aliphatic Amines in Water, J. Am. Chem. Soc. 125, 310–311.

(11) Arnett, E. M., Jones, III, F.M., Taagepera, M., Henderson, W. G., Beauchamp, J. L., Holtz, D., Taft, R. W. (1972) A Complete Thermodynamic Analysis of the “Anomalous Order” of Amine Basicities in Solution, J. Am. Chem. Soc. 94, 4724–4726.

(12) Dai, S., Holt, M. V., Horton, J. R., Woodcock, C. B., Woodcock Patel, A., Zhang, X., Young, N. L., Wilkinson, A. W., Cheng, X., (2020) Characterization of SETD3 methyltransferase–mediated protein methionine methylation, J. Biol. Chem. 295, 10901–10910.

(13) Lewis, Jr., C.A., Jr., Wolfenden, R. (2018) Sulfonium Ion Condensation: the Burden Borne by SAM Synthetase, Biochemistry 57, 3549–3551.

(14) Benson, S. W., Bond Energies, J. Chem. Educ. 1965, 42, 502–518.

(15) Cabani, S., Gianni, P., Mollica, V., Lepor, L. Group Contributions of the Thermodynamic Properties of Non-Ionic Organic Solutes in Dilute Aqueous Solution, J. Solution Chem. 1981, 563–595.

(16) Hall, Jr., H.K., Correlation of the Base Strength of Amines, J. Am. Chem. Soc. 1957, 79, 5441–5444

(17) Mihel, I. Knipe, J.O., Coward, J. K., Schowen, R. (1979) α-Deuterium Isotope Effects and Transition-State Structure in an Intramolecular Model System for Methyl-Transfer Enzymes J. Am. Chem. Soc. 101, 4349–4351.

(18) Iwig, D. F., Booker, S. J. (2004) Insight into the Polar Reactivity of the Onium Chalcogen Analogues of S-Adenosyl-L-methionine, Biochemistry 43, 13496–13509.

(19) Wu, G., McLeish, M. J. (2013) Kinetic and pH Studies on Human Phenylethanolamine N-Methyltransferases, Arch. Biochem. Biophys. 539, 1–8.

(20) Stockbridge, R. and Wolfenden, R. (2009) The Intrinsic Reactivity of ATP and the Catalytic Proficiencies of Kinases Acting on Glucose, N-Acetylglucosamine and Homoserine: a Thermodynamic Analysis. J. Biol. Chem. 184, 22747–22757.

(21) Burgner, Jr., J. W., Ray, Jr., W. J. (1984) On the Origin of the Lactate Dehydrogenase Induced Rate Effect, Biochemistry 23, 3636–3648.

(22) Schroeder, G. K., Wolfenden, R. (2007) The Rate Enhancement Produced by the Ribosome: an Improved Model. Biochemistry 46, 4037–4044.

(23) Radzicka, A., Wolfenden, R. (1995) A Proficient Enzyme, Science 267, 90–93.

(24) Wolfenden, R. (2006) Degrees of Difficulty of Water-Consuming Reactions in the Absence of Enzymes, Chem. Rev. 106, 3379–3396.

(25) Paik, W. K., Lee, H. W., Kim, S. (1975) Non-Enzymatic Methylation of Proteins with S-Adenosyl-L-Methionine, FEBS Letters 58, 39–42.

